# SciCV, the Swiss National Science Foundation’s new CV format

**DOI:** 10.1101/2022.03.16.484596

**Authors:** Michaela Strinzel, Wolfgang Kaltenbrunner, Inge van der Weijden, Martin von Arx, Michael Hill

**Affiliations:** Swiss National Science Foundation, Wildhainweg 3, 3001 Bern, Switzerland; Centre for Science and Technology Studies, Leiden University, PO Box 905, 2300 AX Leiden, The Netherlands

**Keywords:** Research assessment, academic CV, change management

## Abstract

**Background:** The Swiss National Science Foundation (SNSF) tested a new CV format called SciCV to encourage fair, DORA-compliant assessment of grant applicants. It was developed in close collaboration with the academic evaluation community and international experts, introduced through detailed change management and finally tested in its utility by an independent research team of the Center for Science and Technology Studies Leiden (CWTS).

**Methods:** We present the development of the SciCV pilot and its evaluation by the CWTS research group. The analysis comprised both quantitative and qualitative methods, with (i) surveys and semi-structured interviews with applicants and reviewers, (ii) text analysis of the narrative elements of SciCV, and (iii) participant observation in ten evaluation panel meetings.

**Results:** Narrative elements and the inclusion of the academic age were rated as most useful new CV elements, while the inclusion of two metrics, the h-index and the relative citation ratio, were received more critically. The omission of a full publication list had similar numbers of supporters and opponents among applicants and reviewers. Less experienced and junior applicants and reviewers rated the new format generally more positively than more senior applicants and reviewers. The text analysis of narrative elements yielded no significant gender specific differences. The participant observation revealed that the new elements in SciCV broadened the information base used in the evaluation of applicants but did not fundamentally alter traditional, publication-centred evaluation practices.

**Conclusion:** SciCV was a relevant and successful initiative for the SNSF, which showed that the implementation of a new, well-structured CV format is not only feasible but also something that many stakeholders welcome.The extensive experience and results obtained during the change process formed the basis for the development of SciCV 2.0 at the SNSF. It also offers a basis and guidance for other funding organisations planning similar initiatives.

## Introduction

The academic Curriculum Vitae (CV) plays a decisive role at many pivotal points in a researcher’s career, including funding and hiring decisions and the awarding of academic prizes [1–3]. Researchers have a right to fair and transparent CV evaluation and research performing and funding organisations need to ensure that they support the right people for the right reasons. The classic “two-page PDF with publication list” CV often falls short of these requirements. It is primarily list-based and unstructured, making interpretation and comparison difficult, and it emphasises publications over other types of academic achievements and publication quantity over quality. These and other challenges inherent to classic CV formats are well documented [see 4 for a current overview and discussion, 5] and have been prominently highlighted by the San Francisco Declaration on Research Assessment (DORA, 2013), the Leiden Manifesto [6], the Metric Tide [7], the Hong Kong Manifesto [8], and the H-Group [3].

In response to these challenges, various funding organisations have started trialling free-text-based CVs instead, allowing applicants to describe their achievements in their own words, including the Dutch Research Council [9], the Science Foundation Ireland [10], the Luxembourg National Research Fund [11], a group of UK-based funders [12], as well as of course the National Institutes of Health through their well-established biosketch (https://grants.nih.gov/grants/forms/biosketch.html). In 2019, the Swiss National Science Foundation (SNSF) also initiated a pilot project to develop and test a new, standardised CV format called SciCV. Inspired by the ACUMEN Portfolio [13] and the innovative evaluation procedure of the Swiss Science Prize Marcel Benoist [14], SciCV was designed neither as a list-based nor a purely free-text-based CV, but instead aimed at combining the best of both worlds.

For the SNSF, an important aspect of implementing SciCV was change management. In order to achieve better community engagement and acceptance, the SNSF trialled a first version of SciCV in one funding call, after which it reverted back to the original system while evaluating and discussing the results and experiences from the pilot in the community. The SNSF also collaborated with independent, third-party experts to ensure the evaluation of the SciCV project was unbiased. The results of the evaluation and discussions in turn fed into the development of an amended version of SciCV (i.e. SciCV 2.0) which the SNSF plans to introduce for all funding instruments in autumn 2022.

This paper highlights the milestones from the change management process of SciCV, including the development, execution, and professional evaluation of the pilot and the key findings gained from it. It concludes with discussing how these insights informed SciCV 2.0 at the SNSF.

## Methods

A declared aim of the SciCV pilot was to develop it together with international experts and the SNSF’s own research evaluation community to ensure that it would be accepted, supported and adopted by the research community. To best incorporate the various stakeholders’ perspectives, a SciCV change management roadmap was designed consisting of the following four steps:

**Step 1** | Groundwork: In 2018, in multiple online meetings and a two-day workshop in Zurich, Switzerland, the CV Harmonisation-Group (H-Group), consisting of international funders, research infrastructure providers and experts in research on research, discussed various ways to improve academic CVs for fairer research assessment [3].

**Step 2** | Design: In 2019, a working group consisting of representatives from the SNSF’s research evaluation community and administrative offices defined the structure and content of SciCV based on the outputs of the H-group’s work.

**Step 3** | Pilot: In 2020, following a wide-spread communication initiative and the go-live of the SciCV online platform [15], SciCV was mandated for all 346 applications submitted in the April 2020 call in the SNSF Project Funding scheme in biology and medicine. Project Funding is the SNSF’s main funding scheme with a total of approximately 2000 applications per year from all scientific disciplines.

**Step 4** | Analysis and follow-up: In 2020 and 2021, in close collaborations with the Centre for Science and Technology Studies at the University of Leiden (CWTS), extensive analyses of the SciCV pilot and the various participants’ experiences with it were conducted. These experiences served as a basis for improving SciCV.

The evaluation of SciCV in step 4 consisted of four types of data collection and analysis:

### Survey

Two surveys were conducted using the Qualtrics survey platform (05/2020, https://www.qualtrics.com/), one for applicants and one for reviewers. Three types of reviewers were involved in the assessment of proposals for the Project Funding scheme: (i) external reviewers and members of the evaluation panel, consisting of (ii) external panel members and (iii) members of the SNSF’s national research council (they will be referred to as panel members and RC members respectively). The survey asked respondents to rate the usefulness of key elements of SciCV on a scale from 1 (not at all useful) to 5 (extremely useful).Reviewers were also asked to rate the overall usefulness of SciCV in the assessment of applications. Potential differences in the collected responses were analysed with regards to the applicants’ and reviewers’ gender, age, scientific fields within medicine and biology and experience of both applicants and reviewers using the statistical software package SPSS 26 (see table 1 for characteristics of survey respondents).

**Table 1.**
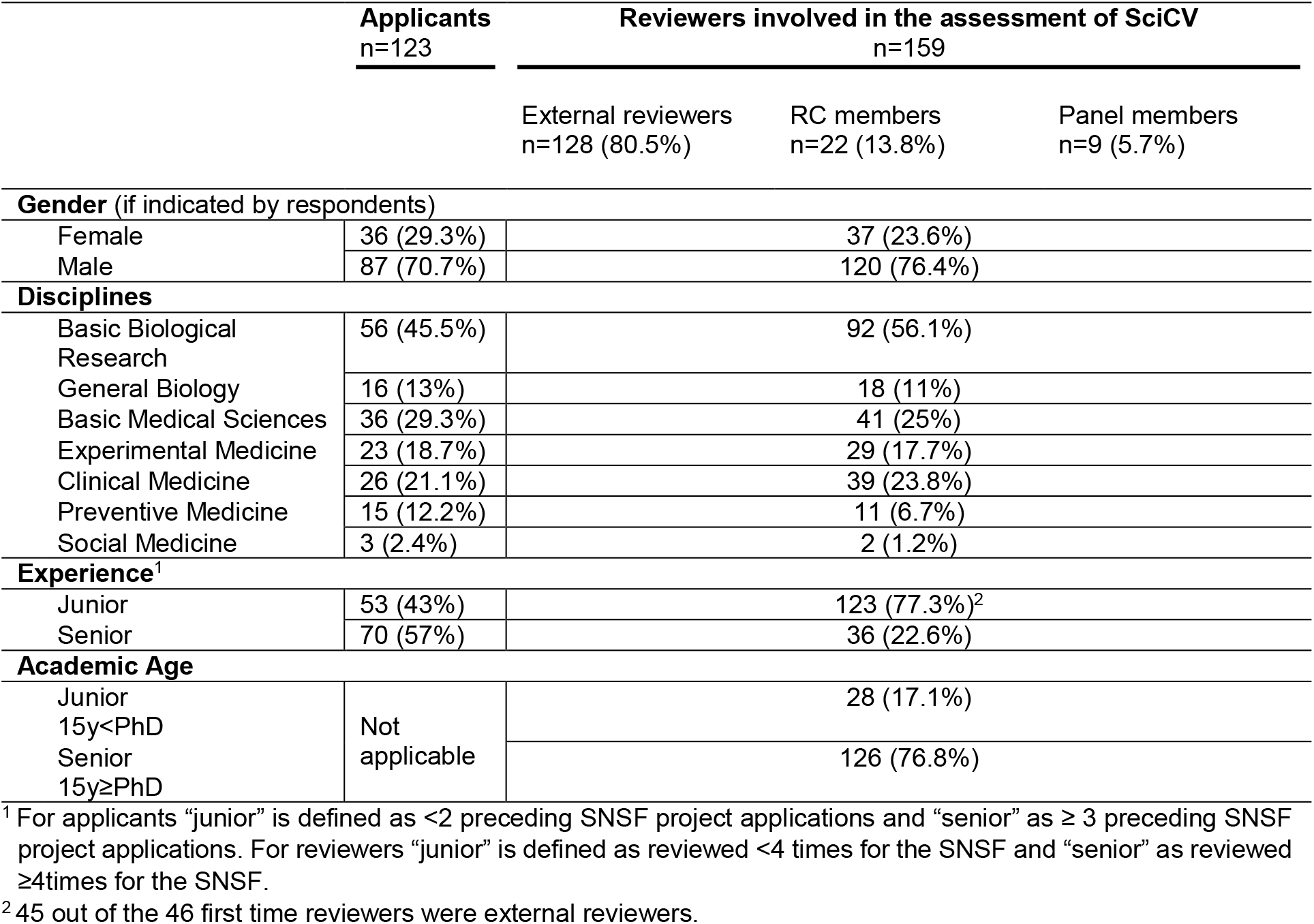
Characteristics of survey respondents.

### Semi-structured Interviews

In-person online interviews with applicants and reviewers were conducted to acquire a more in-depth understanding of the opinions communicated in the survey. Interview partners were recruited after they had indicated their willingness to be contacted in the survey and selected to create a gender balance subset, where possible (see table 2). All interviews were semi-structured. This means that a fixed set of questions was asked but with the possibility for respondents to extensively discuss their responses and potentially raise relevant points not foreseen by the interviewer. In total, 20 interviews were conducted: ten with applicants, four with external reviewers, two with regular panel members, and four with research council members. All interviews were recorded and transcribed verbatim. The interviews were analysed by manually coding them according to their pertinence to the main categories of SciCV.

**Table 2.**
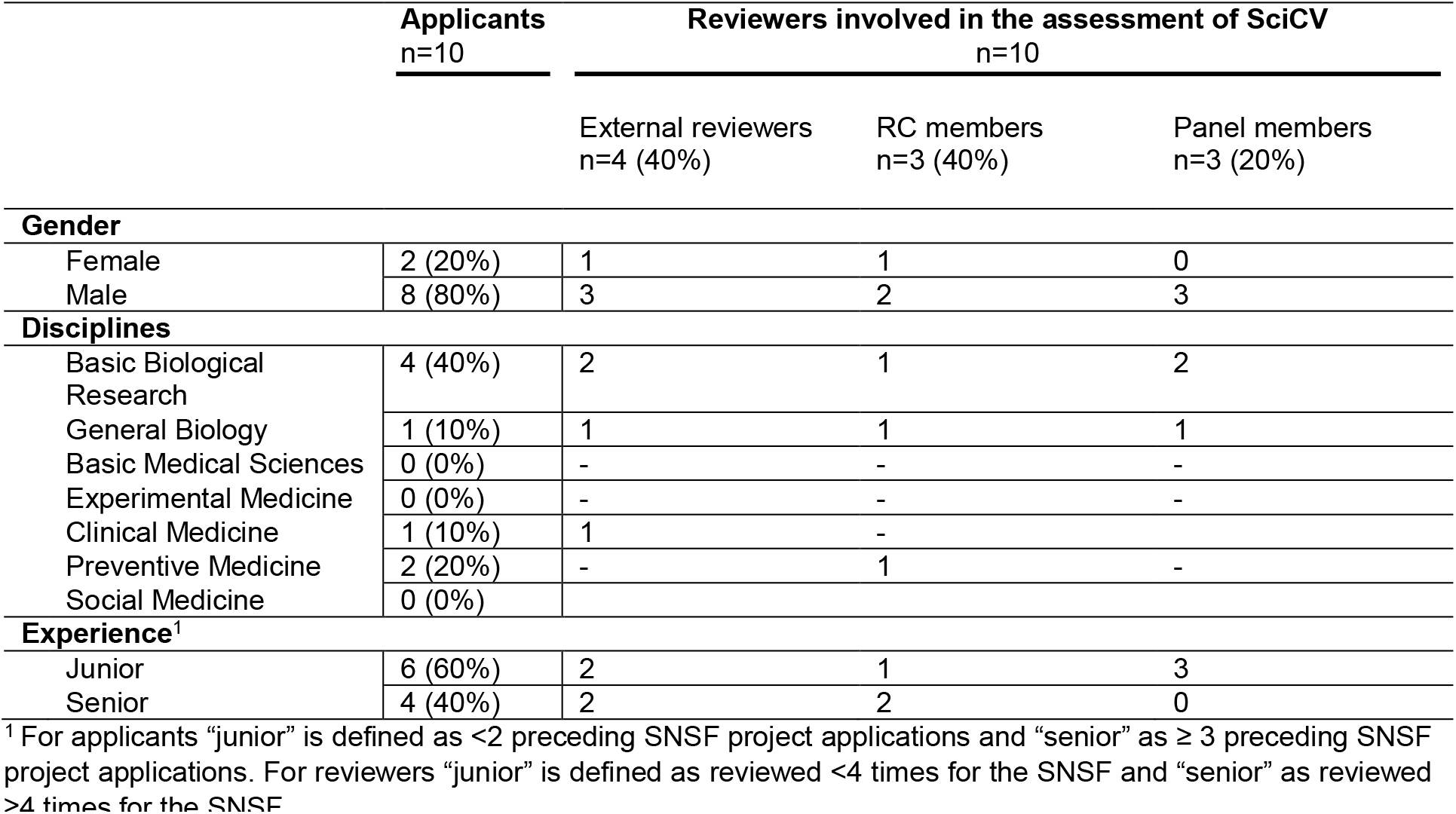
Characteristics of interview partners.

### Text analysis

The narrative elements of SciCV were analysed by means of descriptive statistics of the frequency of pre-defined words to investigate potential gender-specific differences.

Participant observations | Observation data was collected through participant observation in ten separate online evaluation panel meetings. During participant observation, detailed notes on the interaction among the reviewers (e.g. regular panel members and research council members) were taken. In addition to the live attendance, video recordings of the meetings were analysed and the occurrence of certain discussion elements and mentioning of SciCV components were counted systematically.

## Results

In step 2 of the change management roadmap, the working group designed SciCV and structured it into eight sections based also on the outcomes of step 1 [see 3] : (1) Name and Position, (2) Academic Age, (3) H-index, (4) Education / Qualifications, (5) Employment, (6) Funding, (7) Project-related Narrative, (8) Contributions to Science. In order to ensure that SciCV is easy to fill in and machine readable, a dedicated online platform was developed with tight integration of the Open Researcher and Contributor ID (ORCiD, https://orcid.org/).

Section 1 contained the applicant’s ORCiD, their name and position. In section 2 the applicant was asked to provide their academic age (AA), which was defined as the number of full time equivalent (FTE) years of work in academia. The AA had to be calculated from their first academic publication onwards. Additionally, women had to always deduct at least 1.5 FTE years, while men were asked to deduct the actual FTE time during which they were fulfilling child-care duties from their AA for every one of their children born, adopted or otherwise in their responsibility during this time. Section 3 contained the H-index [16] as calculated based upon the Scopus database (https://www.scopus.com/home.uri). Section 4, 5 and 6 consisted of education and qualifications, employment, and funding histories respectively, including information such as dates, institution names and further details like the degree earned, the employment title or the amount of funding acquired, respectively. In section 7 the applicant was asked to write a project-related narrative text, in which they describe how and why they are well-suited to execute the proposed project. This project-related narrative consisted of up to 300 words, any claims made therein had to be substantiated by referencing up to five works (publications, code or any other type of output). Whenever a referenced work was a publication, the SciCV platform automatically pulled the respective publication’s Relative Citation Ratio metric (RCR) [17] from the Dimensions website (https://www.dimensions.ai/), where available and appended it to the citation. Section 8 was similar to section 7 but it consisted of up to four narratives of 200 words and up to four references each, describing the applicant’s most important previous contributions to science. Beyond the maximum of 21 references used in the narrative sections 7 and 8 of SciCV (i.e. five for substantiating the narrative in section 7 and four for each of the four narratives in section 8), no other references could be provided and no complete publication list or similar could be submitted during the application process.

SciCV could be completed on an online platform, which provided eight separate tabs representing the eight sections of SciCV. To use the SciCV platform, applicants had to log in with their ORCiD. Through this log-in process, the applicant’s SciCV was automatically connected to their ORCID account and all information stored in ORCID could easily be pulled into SciCV. If an applicant had a well-maintained ORCID account, this meant that section 1, 3, 4 and 5 as well as all references in section 7 and 8 could be entered into SciCV by selecting the relevant items from a drop-down menu. Filling out section 3 was also facilitated if applicants had an up-to-date Scopus account. Both the ORCID and the Scopus accounts were mandated by the SNSF for applications during this pilot.

SciCV was piloted during the April 2020 Project Funding call in the SNSF Division of Medicine and Biology. This funding call resulted in a total of 346 applications, involving 495 applicants. The applications were first assessed by international, external peer reviewers with one application per reviewer and at least two reviewers per application. Then, drawing on the external peer review reports, applications were discussed in a series of panel meetings taking place online in August 2020. The panels consisted of research council members and other expert participants, who were responsible for determining a ranking of the applications. Ultimately, 129 applications received funding.

### Survey

The survey was part of the analysis of the SciCV pilot conducted by the research group of CWTS Leiden. The applicant survey achieved a response rate of 24,8%, equalling 123 respondents and the reviewer survey a response rate of 12,4% equalling 159 respondents. Figure 1 and figure 2 show the rating scores for key SciCV elements by applicants and reviewers.

**Fig 1.**
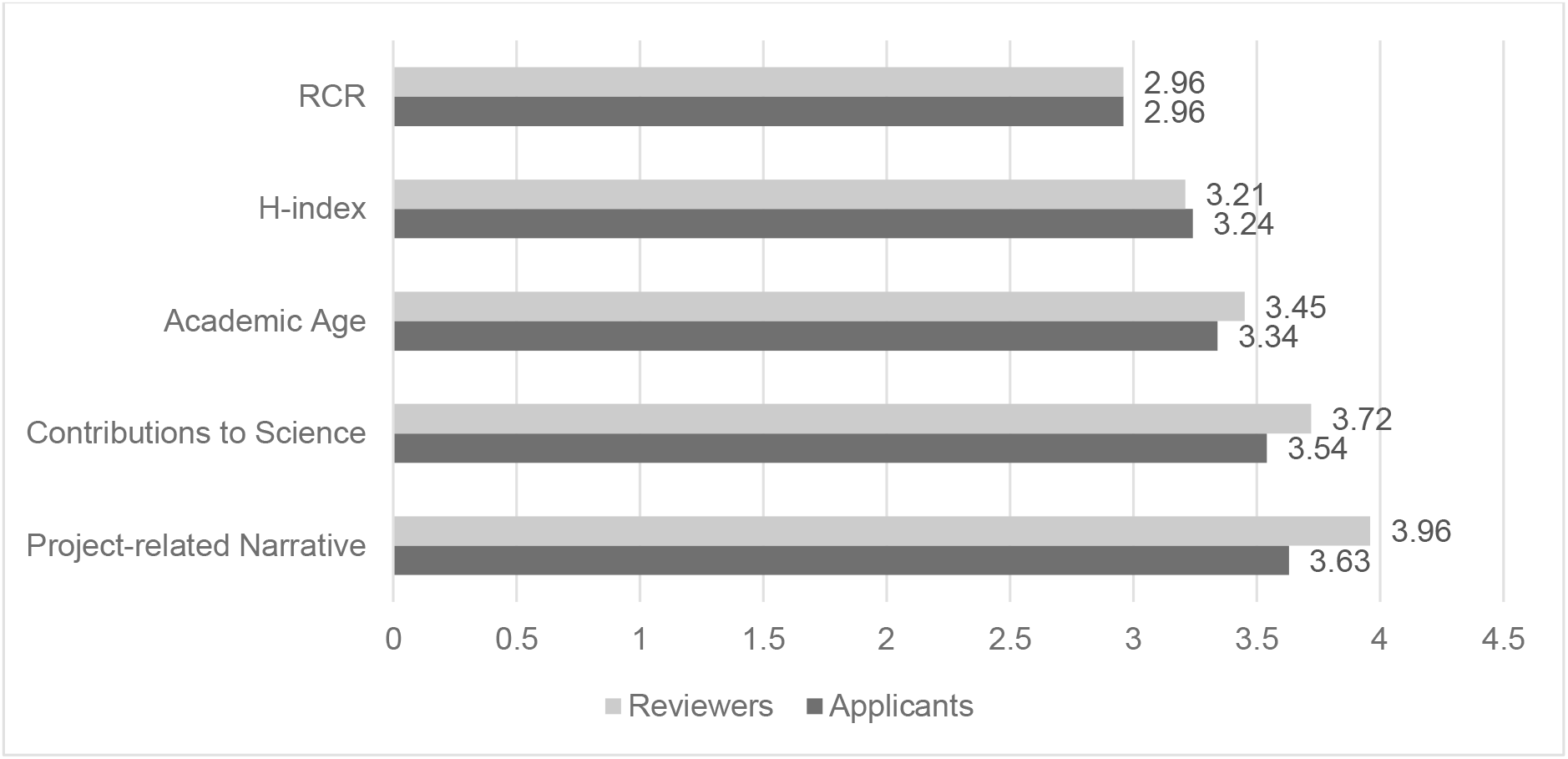
Mean ratings of usefulness per SciCV elements. The figure shows the mean usefulness ratings per SciCV element by applicants and reviewers on a scale from 1 (not at all useful), 2 (slightly useful), 3 (moderately useful), 4 (very useful), to 5 (extremely useful).

**Fig 2.**
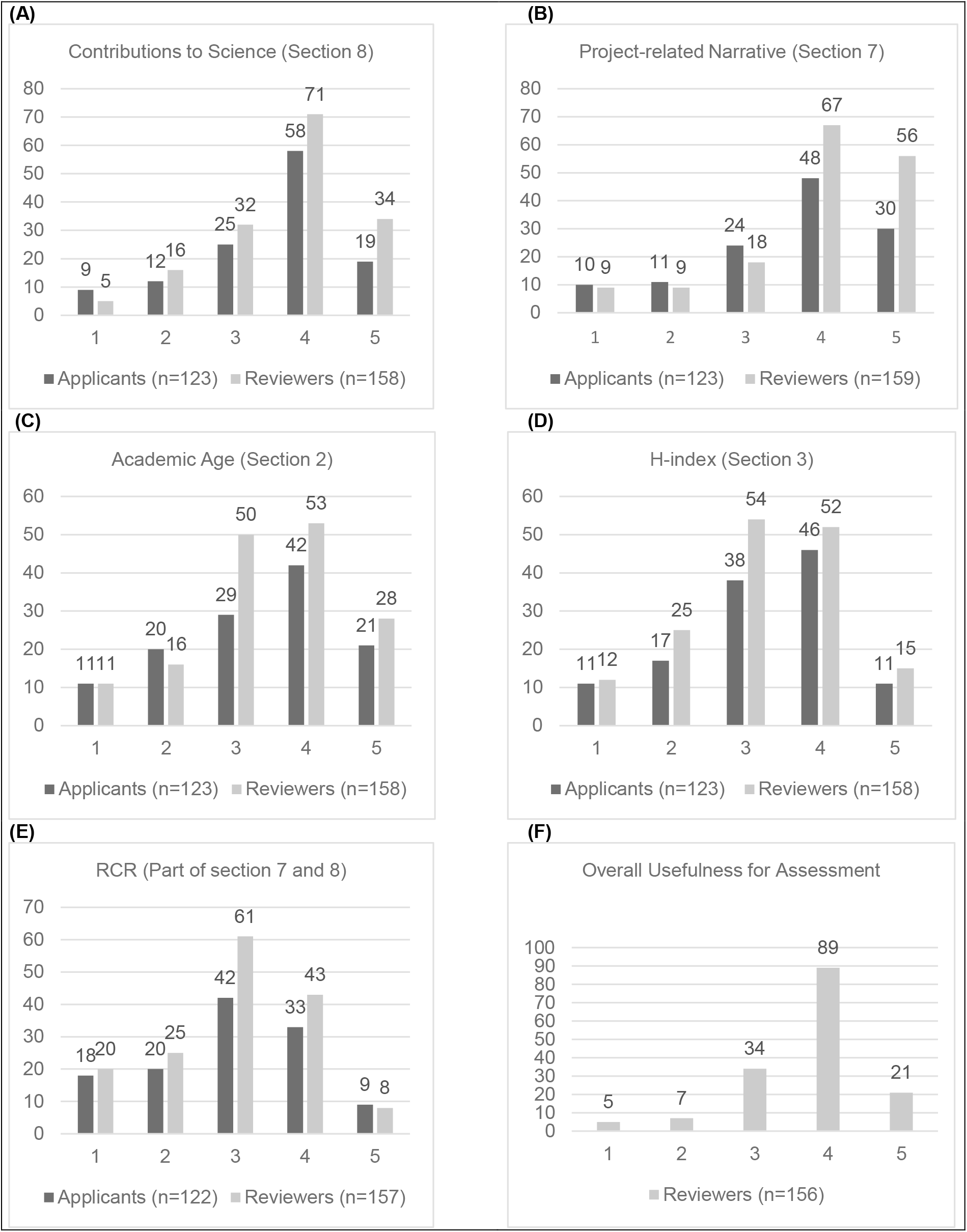
Perceived usefulness of SciCV elements. Figures A-E show the usefulness ratings in total numbers for the single SciCV elements on a scale from 1 (not at all useful), 2 (slightly useful), 3 (moderately useful), 4 (very useful), to 5 (extremely useful) by applicants and reviewers. Figure F shows the raw rating numbers of the usefulness of SciCV for the assessment of applicants by reviewers on the same scale from 1 (not at all useful) to 5 (extremely useful).

Applicants and reviewers alike rated the two types of narratives as most useful among the new elements of SciCV, with the project-related narrative scoring 3.63 (moderately to very useful) out of 5 and the contributions to science 3.54 (moderately to very useful) out of 5 for applicants and 3.96 (very useful) out of 5 for the project-related narrative and 3.72 (very useful) out of 5 for the contributions to science for reviewers (see Fig 2A and B). The inclusion of the academic age in SciCV was equally well received by both applicants and reviewers with a rating of 3.34 (moderately useful) out of 5 for applicants and a rating of 3.45 (moderately useful) out of 5 for reviewers (Fig 2C). The inclusion of the two metrics H-index and RCR were less enthusiastically received by applicants and reviewers but still considered useful with ratings of 2.96 and 3.24 out of 5 (both moderately useful) for the RCR and the H-index for applicants, respectively, and an RCR scoring 2.96 out of 5 and H-index 3.21 out of 5 (both moderately useful) for reviewers (Fig 2D and E). Only the reviewers were asked to rate SciCV’s overall usefulness to assess an applicant. 70% (equalling 110 respondents) of the survey respondents experienced the content of SciCV as very or extremely useful to assess the expertise of the applicants (Fig 2F). The omission of the full publication list was ambivalently received by both applicants and reviewers with roughly equal numbers of advocates and opponents with regard to applicants. 46% of applicants and 43% of reviewers were against omission, 9% of applicants and 10% of reviewers were indifferent, and 45% of applicants and 47% reviewers were in favour (Fig 3).

**Fig 3.**
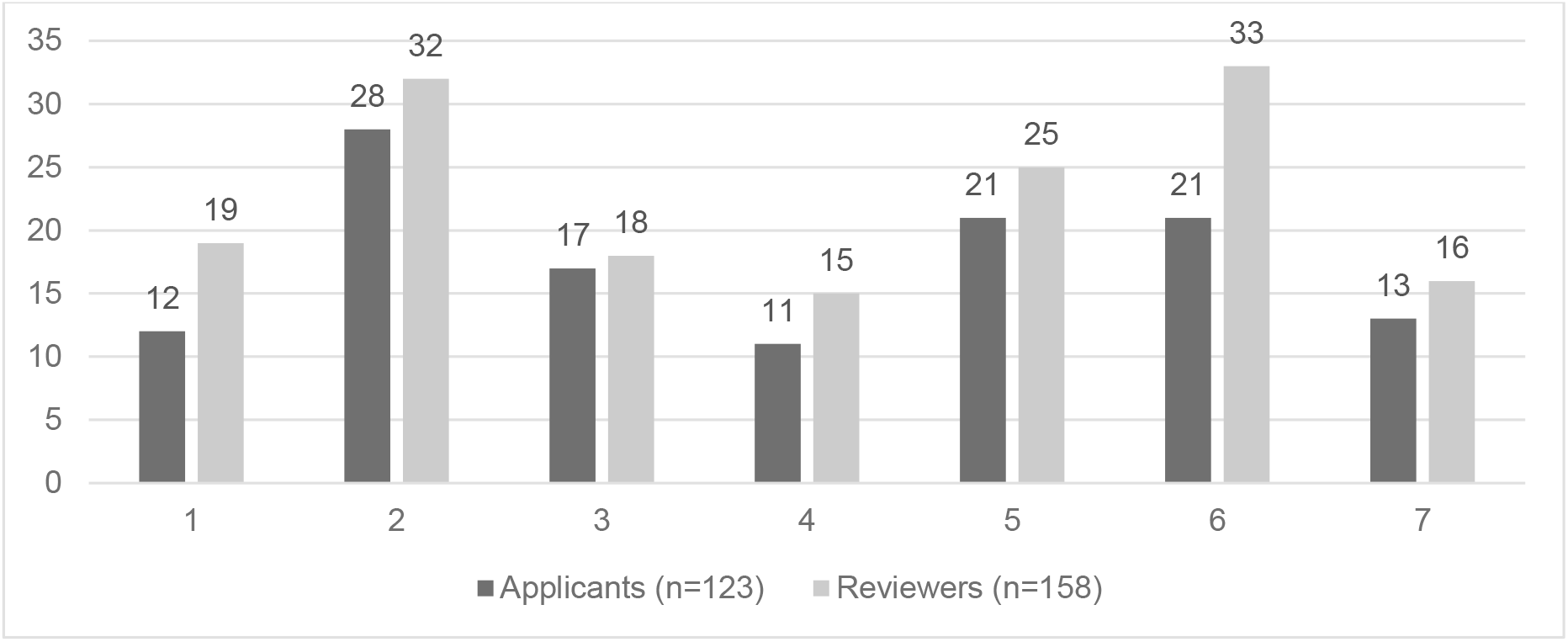
Ratings of agreement with the omission of the full publication list. The figure shows the raw rating numbers of agreement with the omission of the full publication list on a scale from 1 (strongly agree), 2 (agree), 3 (somewhat agree), 4 (neither agree nor disagree), 5 (somewhat disagree), 6 (disagree), to 7 (strongly disagree) by applicants and reviewers.

### Differences in ratings in terms of respondent characteristics

ANOVA analysis showed that the ratings of key novel features of SciCV were influenced by the amount of previous experience that applicants and reviewers had as well as the type of reviewer they are and their experience as reviewers, but not by their discipline or gender (see table 3).

**Table 3.** Differences in mean ratings of SciCV elements depending on respondent characteristics. The table shows the average usefulness ratings of junior and senior applicants and reviewers and the three different reviewer types. Ratings were made on a scale from 1 (not at all useful), 2 (slightly useful), 3 (moderately useful), 4 (very useful), to 5 (extremely useful), except for the “Omission of publication list”, which was made on a scale from scale from 1 (strongly agree), 2 (agree), 3 (somewhat agree), 4 (neither agree nor disagree), 5 (somewhat disagree), 6 (disagree), to 7 (strongly disagree).

| SciCV element | Applicants |  | Reviewers |  |  |  |  |  |
| --- | --- | --- | --- | --- | --- | --- | --- | --- |
|  | Experience<sup>1</sup> |  | Experience<sup>2</sup> |  |  | Type |  |  |
|  | Junior | Senior | 1<sup>st</sup> time | Junior | Senior | External reviewer | Panel member | RC member |
| Project related narrative | 3.87 | 3.44 | 4.13 | 4.05 | 3.53 | 4.14 | 3.56 | 3.05 |
| Contributions to science | 3.79 | 3.34 | 3.72 | 3.79 | 3.56 | 3.86 | 3.33 | 3.05 |
| Academic Age | 3.57 | 3.17 | 3.46 | 3.40 | 3.54 | 3.43 | 3.89 | 3.36 |
| H-index | 3.30 | 3.19 | 3.02 | 3.33 | 3.19 | 3.22 | 3.56 | 3.00 |
| RCR | 2.98 | 2.94 | 2.80 | 3.19 | 2.69 | 2.91 | 3.43 | 2.95 |
| Omission of publication list | 3.91 | 3.97 | 4.41 | 4.13 | 3.19 | 4.27 | 4.33 | 2.32 |
| Overall usefulness for assessment | - | - | 4.02 | 3.78 | 3.26 | 3.93 | 3.67 | 2.64 |

<sup>1</sup> For applicants “junior” is defined as <2 preceding SNSF project applications and “senior” as ≥ 3 preceding SNSF project applications.
<sup>2</sup> For reviewers “junior” is defined as reviewed <4 times for the SNSF and “senior” as reviewed ≥4times for the SNSF.

### Experience

Junior applicants, i.e. having a maximum of two previous SNSF Project Funding applications, were overall more positive about the project-related narrative (average 3.87 vs 3.44; F=3.988 / p=.048), the contribution to science (average 3.79 vs 3.34; F=5.251 / p=.024) and academic age (average 3.57 vs 3.17; F=3.324/ p=.071) than more senior applicants. First time reviewers for the SNSF – all but one of whom were external reviewers -gave significantly higher scores to the usefulness of the project-related narrative compared to more experienced grant evaluators (average 4.13 vs. 3.53; F=3.733 / p=.026) and they agreed more with omission of a complete publication list compared to more experienced reviewers (average 4.41 vs. 3.19; F=4.305 / p=.015). With regard to the overall usefulness of SciCV to assess applicants, which was only rated by reviewers, first time reviewers (average 4.02) had more positive experiences compared to more experienced (mean 3,78 and 3.26; F=8.631 / p=.000) reviewers.

### Type of reviewer

External reviewers and regular panel members gave higher scores to the project-related narrative compared to members of the research council (average 4.14 vs. 3.56 vs. 3.05; F=11.255 / p=.000). A similar picture is found for the contributions to science, which were rated more positively by external reviewers and regular panel members than by members of the research council (average 3.86 vs. 3.33 vs. 3.05; F=7.196 / p=.001). External reviewers and regular members of the evaluation panel agreed with omission of a complete publication list, while members of the research council disagreed (average 4.27 vs. 4.33 vs. 2.32; F=10.273 / p=.000). The overall usefulness to assess applicants based on SciCV was rated more positively by external reviewers and regular panel members compared to members of the research council (average 3.93 vs. 3.67 vs. 2.64; F=27.979 / p=.000).

### Interviews

The interviews provided detailed information on how respondents experienced the practical use of SciCV, allowing understanding in more detail why respondents held certain views and thereby complementing and deepening the overarching trends indicated by the survey.

### Narratives

The interviews indicate that applicants disliked the amount of time needed to author narratives, but also appreciated the opportunity to demonstrate connections between different research activities and highlight achievements for which there were no other dedicated sections. The interviews with reviewers suggest that they appreciated the contextualising aspect of narratives for their evaluation, as they provide an overview of scientific careers and connections that are not discernible from publication lists only. However, redundancy and the use of boastful language in narratives were also criticised by some reviewers.

*“And particularly the project related narrative, I can see that it is a good idea. So okay, to show that you have a clear idea about what you want to do. I mean, it is not something you put together in a few days. So, and it agrees with your research and all these things, so sure I can see why they want it. But if you want to do it properly then this takes time […]”* (Applicant)

*“I think that from a reviewer point of view, especially if you’re slightly tangential to a field, it’s like, it’s really nice to know like, what is the context of this and have the person tell you what they think the context is.[*…*] And I think that what was also useful, as I said, at this particular review I did go in and read through some of the papers. And I could see that then, it really did give me context. And of course like, this person is presenting themselves in the best possible way. So, it did help bias me to like the paper, to be honest. But that being said, I think that it is helpful. And I think it’s also helpful to see threads between papers. And so like, this is sort of like what this person is attempting to do. And then especially if that then is related to the grant, it’s super valuable. So, I actually really like that as a reviewer.[…]”* (External reviewer)

### Academic Age

Some applicants and reviewers disagreed with the way AA was calculated, for example the fact that counting started with the year of the first publication rather than the year of graduation and that different types of parental leave were factored into the formula for men and women. While men could only deduct the effective time spent on childcare duties, women had to deduct at least 1.5 FTE per child to take into account restrictions beyond the actual absence during maternity leave, such as possible restrictions during pregnancy and breastfeeding, limited mobility, and flexibility etc. In some cases, reviewers also voiced distrust in some of the AA values calculated by applicants.

*“And then it ended up like out of my academic career which was eleven years, I got eight [years]. Which is clearly an advantage. But then, I mean I will be honest with you, if I have had taken one point half years for each one of my children, my career would have ended before it has even started. (…) Something has to be done, but I don’t think this is the good way.”* (Applicant)

*“[academic age] was very often really wrong. You had a 50 year old man, who has been working in research institutions, often in leading positions, since 30 years, who put the academic age of 5 years (…) I mean they’ve got a calculator and I did something, but then there’s still details, still he has been working, he has had this and this grants, he has been working for 5 years as postdoc there and then as a research leader there, and as a director of a university, doing research or an institute there and then the number of years is so much inferior. It’s ridiculous. Then either the CV is wrong, like the previous jobs or the number is wrong.”* (Research council member)

### Metrics

The interviews with reviewers and applicants suggest that the H-index plays an important role in terms of providing a minimum threshold. The index becomes a problem when it is particularly low, but a high index is never a determinant in a competition between applicants. There was moreover widespread awareness of the limitations of the H-index by applicants and reviewers. In contrast, most applicants and reviewers were not familiar with the RCR and how exactly it is calculated. Those who understood the metric appreciated that it is not age biased (in comparison to the H-index) and that it aims to compare papers with similar papers, rather than using the journal reputation as a proxy for the quality of the article (as the journal impact factor does).

*“For me it was actually the first time in this format that I discovered [the RCR]. I think it is useful, I guess the only problem is a bit that it is this specific*… *I guess it is not possible otherwise. Because it only calculates the RCR for publications that are listed for more than at least two years.[…] a lot of my main publications I couldn’t even indicate an RCR because it is not available yet. That is, especially for younger people (*…*) like if you are doing a grant application quite early in your academic career, that can be a problem. Because perhaps a lot publications are quite recent.”* (Applicant)

*“Also things like H-index are really again biased towards older people. So, your H-index can only be as many papers as you have. So, if you’re early in your career, your H-index is inherently going to be lower. And I think that the new indicator [the RCR] (…) at least tries that to address by field. Because like, in my field, papers take, a big paper would take two years at least for a post-doc. Whereas like there are other fields that are just much faster, because the whole process is faster.”* (External reviewer)

### Omission of the publication list

Applicants and reviewers in support of the omission felt that the sample of relevant publications highlighted by the applicant in the narrative elements is sufficient for evaluation and that the omission brings benefits in restricting certain biases. Opponents of the omission feared an incomplete picture of a researcher’s profile as well as a lack of verification of claims made in the narratives.

*“If I think about somebody who reviews [an applicant’s CV] - especially if they are little bit, let’s say, more experienced researcher, sometimes [the length of a full publication list] gets too excessive, right. And I wouldn’t even look at it. I think for somebody at my position, it is sometimes nice if you can refer to work that you have done before. I mean even if it’s longer than five years. But I can understand it, I mean at the same time, yeah, if you have a 20 pages list of publications, I wouldn’t even look at it, to be a hundred percent honest.” (Applicant)*

*“Obviously, you are sensitive to journal types and names. It’s almost inevitable that you look at that. I think […] automatically your brain is trying to find quick determiners, right. So, whether you tell yourself not to bias yourself towards it, it’s inevitable. Because you are sensitive to these measures and metrics. So, that is why I am - and now we’re already talking about SciCV - this is why I really encourage these types of changes and go more towards bio sketches. Because if I have to review for NIH or something else, then I do definitely look at the bio sketch. I think that’s very important. Yes.”* (Regular panel member)

### Text analysis of all SciCVs

The narrative elements of the SciCV pilot were analysed to understand to what extent men and women present themselves differently. Here only the results from section 8 (i.e. up to four contributions to science per applicant) with regard to gender differences are presented, as they provide a more comprehensive data set in terms of volume than the project-related narrative of section 7 (1980 contributions to science vs. 495 project-related narratives).

To analyse potential differences in the self-description of male and female applicants, descriptive statistics on word frequencies were employed. Words included in the analysis were predefined terms, which are associated with an assertive style of self-representation such as ‘success’, ‘discovery’, ‘expert’, ‘novel’, ‘pioneer’ and terms related to successful publishing such as ‘paper’, ‘publication’, ‘cited’, ‘citations’ (see table 4 for all predefined terms). Wherever the occurrence ratio of these terms was significantly higher or lower than the proportion of women in the overall population (i.e. 28%), a trend (however weak) was inferred. An additional important value was the relative occurrence of terms in percentage of all contributions submitted by men and women respectively – which allows to estimate how representative particular results are for their gender overall.

**Table 4.** Occurrences of pre-defined terms per gender in the section “contributions to science”.

| Terms | Total occurrence | Total occurrence male/female applicants | Occurrence female applicants in % | % occurrence among female applicants | % occurrence among male applicants |
| --- | --- | --- | --- | --- | --- |
| the first | 302 | 204/98 | 32.45 | 18.01 | 14.21 |
| novel | 212 | 155/57 | 26.887 | 10.48 | 10.794 |
| publication | 156 | 102/54 | 34.615 | 9.926 | 7.103 |
| paper | 113 | 82/31 | 27.434 | 5.699 | 5.710 |
| discovered | 89 | 64/25 | 28.09 | 4.596 | 4.457 |
| success | 85 | 54/31 | 36.471 | 5.699 | 3.76 |
| author | 85 | 60/25 | 29.412 | 4.596 | 4.178 |
| discovered | 89 | 64/25 | 28.09 | 4.596 | 4.457 |
| discovery | 80 | 65/15 | 18.75 | 2.757 | 4.526 |
| expert | 79 | 46/33 | 41.772 | 6.066 | 3.203 |
| article | 77 | 53/24 | 31.169 | 4.412 | 3.691 |
| independent | 71 | 50/21 | 29.577 | 3.86 | 3.482 |
| grant | 68 | 45/23 | 33.824 | 4.228 | 3.134 |
| innovative | 54 | 36/18 | 33.333 | 3.309 | 2.507 |
| pioneer | 54 | 38/16 | 29.63 | 2.941 | 2.646 |
| unique | 48 | 34/14 | 29.167 | 2.574 | 2.368 |
| award | 46 | 24/22 | 47.826 | 4.044 | 1.671 |
| expertise | 45 | 28/17 | 37.778 | 3.125 | 1.95 |
| cited | 43 | 35/8 | 18.605 | 1.471 | 2.437 |
| promising | 40 | 25/15 | 37.5 | 2.757 | 1.741 |
| achievement | 36 | 27/9 | 25 | 1.654 | 1.880 |
| major contribution | 25 | 18/7 | 28 | 1.287 | 1.253 |
| fellowship | 20 | 12/8 | 40 | 1.471 | <1% |
| prestigious | 16 | 11/5 | 31.25 | <1% | <1% |
| excellent | 14 | 11/3 | 21.429 | <1% | <1% |
| citations | 12 | 11/1 | 8.3333 | <1% | <1% |
| breakthrough | 6 | 6/0 | NA | NA | NA |
| leadership | 5 | 3/2 | 40 | <1% | <1% |
| Nature (journal) | 19 | 15/4 | <1% | <1% | 1.045 |
| Science (journal) | 5 | 4/1 | <1% | <1% | <1% |

The analysis showed only minor differences between male and female applicants in their narratives. Women overall were slightly more likely to use the terms “publication” (34,6%) and “article” (31,1%) compared to men (relative occurrences in the contributions of the respective gender 9.9% vs 7.1%; 4.4% vs. 3.69%), “grant[s]” (47.8%), “award[s]” (40%), and “fellowship[s]” (33.8%) (relative occurrences in the contributions of the respective gender 4.2% vs. 3.13%; 4% vs. 1.6%; 1.4% vs. 0.83%) as well as terms highlighting their expertise, e.g. “the first” (32.45%), “expert” (41.7%), “innovative” (33.3%) (relative occurrences in the contributions of the respective gender 18% vs. 14.2%; 6.1% vs. 3.2%; 3.3% vs. 2.5%). Note, however, that the overall occurrences of these terms are very low, with the most frequently used term “publication” appearing only 156 times in 1980 contributions. On the other hand, women were less likely to indicate the citedness of their publications; either in terms of specific numbers of “citations” (8.3%) or in terms of them being highly “cited” (18.6%). Again, however, the occurrences of these citation-related terms are very low, with the more frequently used term “cited” appearing only 43 times in 1980 contributions).

### Participant observation

The data collected through participant observation in the ten panel meetings provided insights into how reviewers practically handled SciCV in the evaluation and how the format mediated the decision-making process under time and resource constraints. An overarching trend was that discussions in the evaluation panels focused on the quality of the research proposals. Reviewers spent comparatively less time discussing the applicants’ CVs, and there were no cases where reviewers fundamentally disagreed about the overall quality of the CVs. It is important to note, however, that 31% of applications were evaluated via a “triage”-mechanism, which is a shortened procedure for applications far from the funding line. In general, triaged applications are not discussed in detail during the evaluation meeting unless a panel member asks to do so.

From the discussions among reviewers during the panel meetings, it became clear that SciCV did in fact not fundamentally alter widely established evaluation practices for CVs. Publication-centric criteria remained prominent in the assessment of the quality of applicants’ CVs and their perceived ability to carry out the proposed project. Reviewers focused on formal achievements such as a quantity of output, citations to papers, publications in reputed journals, and funding track records.

Despite clear instructions to use only the information provided in SciCV to assess applicants, in many cases online sources were nonetheless consulted by panel members and research council members to retrieve additional information not contained in SciCV (e.g. PubMed, Google Scholar or personal / institutional website of applicants). In particular, full publication lists, which were not part of SciCV, played an important role in the evaluation work of panel members and research council members, being explicitly mentioned in 37% of all applications discussed (see table 5). For 10% of all applications discussed, reviewers pointed out publications in particularly journals such as Nature, Science, or Cell as evidence of success. In 16% of all cases, reviewers looked up biological age to put applicants’ achievements into perspective.

**Table 5.** Observations panel meetings.

|  | **N=240 applications discussed**<br>**(triage cases excluded)** | **Mentioned in % of cases** |
| --- | --- | --- |
| Biological age | 39 | 16% |
| Publication list/record | 89 | 37% |
| Funding track record | 38 | 16% |
| PubMed profile/similar sources | 2 | 1% |
| Personal website of applicant | 4 | 2% |
| Journal names | 24 | 10% |

## Discussion

SciCV is a new CV format of the SNSF, which was developed, trialed and introduced in a systematic change management process. The rationale of the standardised CV format including text-based elements was to overcome challenges associated with the classic unstructured and list-based academic CV. The change process included the development of a first SciCV version, its piloting in one call of the Project Funding scheme of Division Biology and Medicine, and a subsequent evaluation by an independent research group of CWTS Leiden. Drawing on the results and experience made with the SciCV pilot and following community consultations, the SNSF developed an adapted version, SciCV 2.0.

The change management process related to the SciCV pilot was a relevant and successful initiative at the SNSF. Piloting a new CV format in a real-life setting allowed us to identify its strengths and weaknesses and to collect data from applicants as well as reviewers as a basis for further improvement of the CV format in a subsequent step. The participation of the Swiss research community in the process not only led to important discussions on evaluation practices and research culture, but also increased the acceptance of SciCV. The results of the extensive analysis by the research group of CWTS Leiden showed that SciCV as a whole was well received by applicants and reviewers and that most stakeholders saw value in this new CV format. Applicants and reviewers alike welcomed the narrative elements and the inclusion of the academic age in SciCV. Other aspects, however, especially the metrics and the omission of the publication list, were perceived more critically. The experience of the applicants and reviewers was a decisive factor in the ratings of the new format’s usefulness. Younger, less experienced applicants and reviewers were generally more open to the CV format changes than those, who were probably more used to the classic “two-page PDF with publication list” CV. This finding suggests that the perceived usefulness of key features is to some extent a question of habit and that new elements are likely to be more favourably seen as applicants and reviewers become more used to them. Despite their acknowledged value in the evaluation, there were concerns about gender-specific differences in the free-text elements, which were partly confirmed by previous findings [18]. While the analysis did not find such differences, the SciCV pilot did not provide a substantial enough data set to conclusively deny or confirm any such gender effects. Absence of evidence is of course not evidence of absence, therefore continued monitoring of potential biases in free-text elements will be important in the future.

Not surprisingly, SciCV alone had only a limited effect on the adherence to DORA-conformity during evaluation. Although the new CV elements did broaden the basis of evaluation in this pilot, reviewers still heavily relied on traditional, publication-oriented indicators. Other funding organisations experimenting with new, text-based CV format such as the Science Foundation Ireland report similar findings [10], highlighting the need for additional accompanying measures such as clear guidelines or training.

## Outlook

SciCV 2.0 was developed drawing upon the discussion, experiences and results generated during the change process. The adapted format still combines free-text and list-based elements. It includes those elements of the SciCV pilot, which were highly rated, including the narratives and the academic age, while leaving out less positively rated elements such as metrics. SciCV 2.0 is planned to be implemented across the SNSF as of autumn 2022. The SNSF plans continues monitoring to be able to incrementally improve SciCV.

